# Mapping the human kinome in response to DNA damage

**DOI:** 10.1101/385344

**Authors:** Michel Owusu, Peter Bannauer, Athanasios Mourikis, Alistair Jones, Joana Ferreira da Silva, Michael Caldera, Marc Wiedner, Charles-Hugues Lardeau, Jörg Menche, Stefan Kubicek, Francesca Ciccarelli, Joanna I. Loizou

## Abstract

We provide a catalog for the effects of the human kinome on cell survival in response to DNA damaging agents, selected to cover all major DNA repair pathways. By treating 313 kinase-deficient cell lines with ten diverse DNA damaging agents, including seven commonly used chemotherapeutics, we were able to identify kinase specific vulnerabilities and resistances. In order to identify novel synthetic lethal interactions, we investigate the cellular response to carmustine for 25 cell lines, by establishing a phenotypic FACS assay designed to mechanistically investigate and validate gene-drug interactions. We show apoptosis, cell cycle, DNA damage and proliferation after alkylation or crosslink-induced damage for selected cell lines and rescue the cellular sensitivity of DYRK4, EPHB6, MARK3, PNCK as a proof of principle for our study. Our data suggest that some cancers with inactivated DYRK4, EPHB6, MARK3 or PNCK gene could be particularly vulnerable to treatment by alkylating chemotherapeutic agents carmustine or temozolomide.

## Introduction

The DNA damage response (DDR) is elicited by a complex and far-reaching network of proteins that are commonly deregulated in human pathologies, including cancer^1^. Besides surgery, the most common treatment for cancer patients is radio- or chemotherapy. To date, some of the commonly used chemotherapeutic compounds are DNA damaging agents^2^. DNA damaging agents can cause cell death by targeting either DNA directly or proteins implicated in DNA repair and replication, cell cycle regulators or signal transducers. Protein kinases are an important group of signal transducers and are often deregulated in human cancer^3^, making them particularly interesting to study within the signaling context of the DNA damage response. Moreover, kinases represent an important group of drug targets^4^, due to their enzymatic function, and therefore results from loss-of-function studies with kinases are more likely to be translated into a therapeutic setting. Kinases have broad functions following DNA damage. For instance, the ATM superfamily which includes ATM, ATR and DNA-PKcs (encoded by the gene PRKDC) is involved in sensing or amplifying initial signals of DNA lesions^5,6^. CHK1 and CHK2 kinases, regulate cell cycle progression in response to DNA damage, providing time for DNA repair^7,8^. Other kinases such as ABL1 are involved in transducing or fine tuning signals resulting from DNA damage, which can ultimately lead to survival, senescence or cell death^9^. Though some kinases have been studied in depth, the role of many is still not known^10^. Here, we used CRISPR-Cas9 to individually delete expressed and non-essential kinases in human HAP1 cells. Next, we performed a drug screen using DNA damaging agents, selected to cover all major DNA repair pathways, to map drug specific sensitivities and resistances. We validated selected drug-gene interactions in response to alkylation-induced damage and assessed the contribution of apoptosis, DNA damage, cell cycle arrest and proliferation leading to cellular sensitivities or resistances by designing and utilizing a phenotypic assay.

## Results

We used CRISPR-Cas9 to target 313 expressed and non-essential kinases in human HAP1 cells and produce clonal knock-out cell lines^11^ (**Supp. Table 1**). The kinase genes targeted cover all groups, according to the standard classification scheme of kinases^12^, hence ensuring coverage (Figure 1a). To examine the response of the non-essential human kinome to a broad range of DNA damaging agents, we first designed and optimized our approach using DNA repair deficient cell lines, where we were able to recover known gene-drug interactions (Figure 1b & Supp. Figure 1). Based on this approach, we selected 10 compounds that (1) induce different types of DNA damage and thus utilize distinct DNA repair pathways and (2) are frequently used as chemotherapeutics (Figure 1c). Next, we exposed the 313 kinase-deficient cell lines to these compounds at four concentrations and assessed cellular survival after three days (Figure 1b).

**Figure 1.**
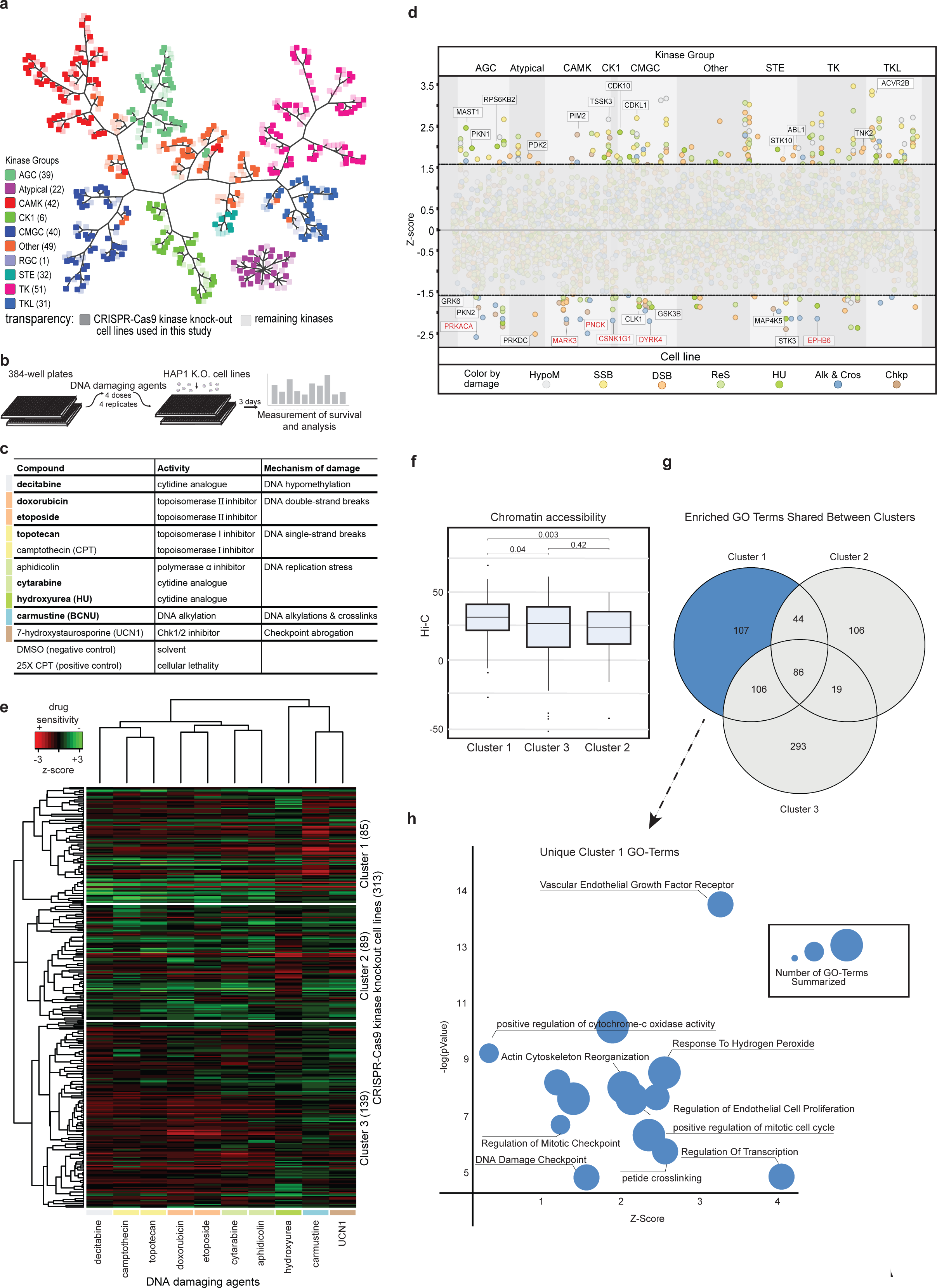
Survival of non-essential kinome in response to DNA damage. (a) Kinome tree representing all kinases^12^. In bold squares are the kinases targeted by CRISPR-Cas9 (313) and in light squares are the remaining kinases. (b) Workflow of survival assay. Dose responses were performed with 4 concentrations in 4 replicates. Cells were incubated with compounds for 3 days and survival was performed using a luminescent readout. (c) List of the 10 DNA damaging compounds selected for use in the survival assay. Compounds with similar modes of action share the same color label. (d) Survival response of cell lines to compounds: kinases from the same kinase groups are clustered together in columns. Each vertical line represents a particular knockout cell line with all of its gene-drug interactions. Compounds are depicted as different color-coded bubbles. HypoM (hypomethylating agent, Decitabine), SSB (single-strand break inducing agents), DSB (double-strand break inducing agents), ReS (replication stress inducing agents), HU (replication stress inducing agent hydroxyurea), Alk & Cros (alkylating and crosslinking agent, BNCU), Chkp (Chk1 inhibitor). Z-scores were calculated for the area under the curve (AUC) of 4 concentrations across the mean of 4 replicates, for each cell line. Lines are set at z-scores greater than 1.65 or less than −1.65 (p<0.05). The names of some expected or known interactions are labeled in black font. Names in red font are examples of lethal interactions after carmustine treatment. AGC= protein kinase families A, G and C; CAMK= Calmodulin/Calcium regulated kinases; CK1= Casein kinase 1; CMGC= CDK, MAPK, GSK3, CLK family; RGC= Receptor guanylate cyclases; STE= STE7, STE11 and STE20 homologs; TK= Tyrosine kinases, TKL= Tyrosine kinase like. (e) Clustering of 313 kinase-deficient cell lines in response to diverse DNA damaging agents reveals three distinct clusters (left dendrogram): Cluster 1 is characterized by a sensitivity to carmustine, Cluster 2 by a sensitivity to hydroxyurea and Cluster 3 by a sensitivity to DNA double-strand break inducing agents, notably etoposide and doxorubicin. Compounds with similar modes of action (color labels) are closer in neighborhood (top dendrogram): topoisomerase II inhibitors (doxorubicin, etoposide), topoisomerase I inhibitors (topotecan, camptothecin), replication stress inducing agents by fork staling (cytarabine, aphidicolin). (f) Difference in chromatin accessibility of genes from Cluster 1, defined by sensitivity to carmustine compared to genes from Cluster 2 or 3. Increasing Hi-C values correspond to increasing chromatin accessibility. (g) Gene ontology (GO) term enrichment analysis for Clusters 1 - 3. (h) GO terms enriched for Cluster 1 uniqely.

Based on literature, most cell lines showing strong sensitivity or resistance to the compounds were anticipated (Figure 1d, **Supp. Table 1**). For instance, PRKDC depleted cells showed the strongest sensitivity to DNA double-strand break inducing agents, etoposide and doxorubicin, whereas ABL1 depleted cells showed resistance those agents^9^.

A clustering of the cell lines by their sensitivity to the 10 compounds revealed 3 clusters, characterized by high sensitivity to carmustine (Cluster 1), hydroxyurea (Cluster 2) and DNA double-strand break inducing agents, such as etoposide and doxorubicin (Cluster 3) (Figure 1e, **Supp. Table 1**). As expected, we found that compounds with similar modes of action were closer in the clustering, as illustrated by topoisomerase II inhibitors (doxorubicin and etoposide), topoisomerase I inhibitors (topotecan and camptothecin) and agents that induced replicative stress (aphidicolin and cytarabine). Due to their clustering with DNA double-strand break inducing agents, our data support the notion that DNA double-strand breaks, following replication fork stalling and collapse, are one of the primary sources of cellular death after treatment with lethal concentrations of aphidicolin, cytarabine or topoisomerase I inhibitors^13^ (Figure 1e). Interestingly, Cluster 1 was significantly enriched for genes associated with increased chromatin accessibility, compared to clusters 2 and 3 (Figure 1f). Alkylating agents, carmustine and temozolomide (TMZ), have been reported to have a global effect on nuclear organization and chromatin structure, inducing chromatin condensation and gene silencing^14^. We therefore reasoned that in the absence of a kinase in Cluster 1, cellular death upon carmustine treatment may be due to an alkylation induced synthetic lethality. Differential gene ontology (GO) term enrichment analysis (Figure 1g, **Supp. Table 2**) confirmed that Cluster 1 was uniquely enriched for terms previously associated to cellular response to alkylating or crosslinking agents: upregulation of vascular endothelial growth factor receptors^15^, induction of oxidative stress^16^ (cellular response to hydrogen peroxide, positive regulation of cytochrome-c oxidase activity), which in turn leads to actin cytoskeleton reorganization^17^ (Figure 1h).

Since kinases are highly associated with cancer^18^, and moreover being enzymes they are potentially amenable to chemical inhibition, we focused on carmustine dependent synthetic lethal interactions. Moreover, we found that cell lines lacking MARK3, PRKACA, CSNK1G1, PNCK, DYRK4 or EPHB6 in combination with carmustine, showed the strongest unreported synthetic lethal interaction in the screen (Figure 1d). To validate and further dissect the mechanism of cellular sensitivity to the drug, we measured DNA damage, apoptosis, cell cycle phases and proliferation in those cell lines in a FACS-based phenotypic assay (Figure 2a) with the markers γH2AX, TUNEL, DAPI and EdU, respectively. As controls, we included cell lines lacking proteins previously linked to the signaling of DNA damage or DNA repair (PRKDC, ABL1^19^, PDK2^20^, PIM2^21^ and TNK2^22^), cell cycle (CDK10, CLK1 and CDKL1), cell death (GRK6^23,24^, GSK3B^25^, MAST1^26^, STK10^27^ and STK3^28^) and a gene with a strong general resistance to DNA damaging agents as revealed in our study (TSSK3) (Figure 1h). Carmustine is a chemotherapeutic agent used for the treatment of several types of cancers, particularly those relating to the nervous system, such as glioblastoma^29,30^. It is a bifunctional alkylating agent that produces DNA mono-alkylation adducts as well as DNA intra- and interstrand crosslinks (ICLs)^31,32^. Almost all of the lesions (90 - 95%) produced by bifunctional alkylating agents are mono alkylation adducts^33^, such as N^7^-methylguanine or O^6^-methylguanine. However, the less abundant (ca. 5%) DNA crosslinks, particularly ICLs, form the most deleterious lesions^33^ and can interfere with replication or transcription and trigger apoptosis and cell cycle arrest^32^. In order to confirm that the predominant cause of cellular toxicity to carmustine was due to the effects of alkylation, we used the monofunctional alkylating agent temozolomide (TMZ), which does not produce crosslinks and is often used as a superior replacement therapy to carmustine^34^, and the crosslinking agent oxaliplatin (Supp. Figure 1b). We chose concentrations of the compounds that moderately affect wild-type cells and as expected, both compounds induced apoptosis, DNA damage, G2/M cell cycle arrest and a reduction of proliferating cells in a dose- and time- dependent manner^35,36^ (Figure 2b-c). Hierarchical clustering of the cell lines according to apoptosis confirmed that the cellular sensitivity to carmustine was predominantly due to alkylation induced synthetic lethality: the most sensitive survival interactions after carmustine (Figure 1h) showed the highest apoptosis after TMZ in the phenotypic assay (Figure 2b).

**Figure 2.**
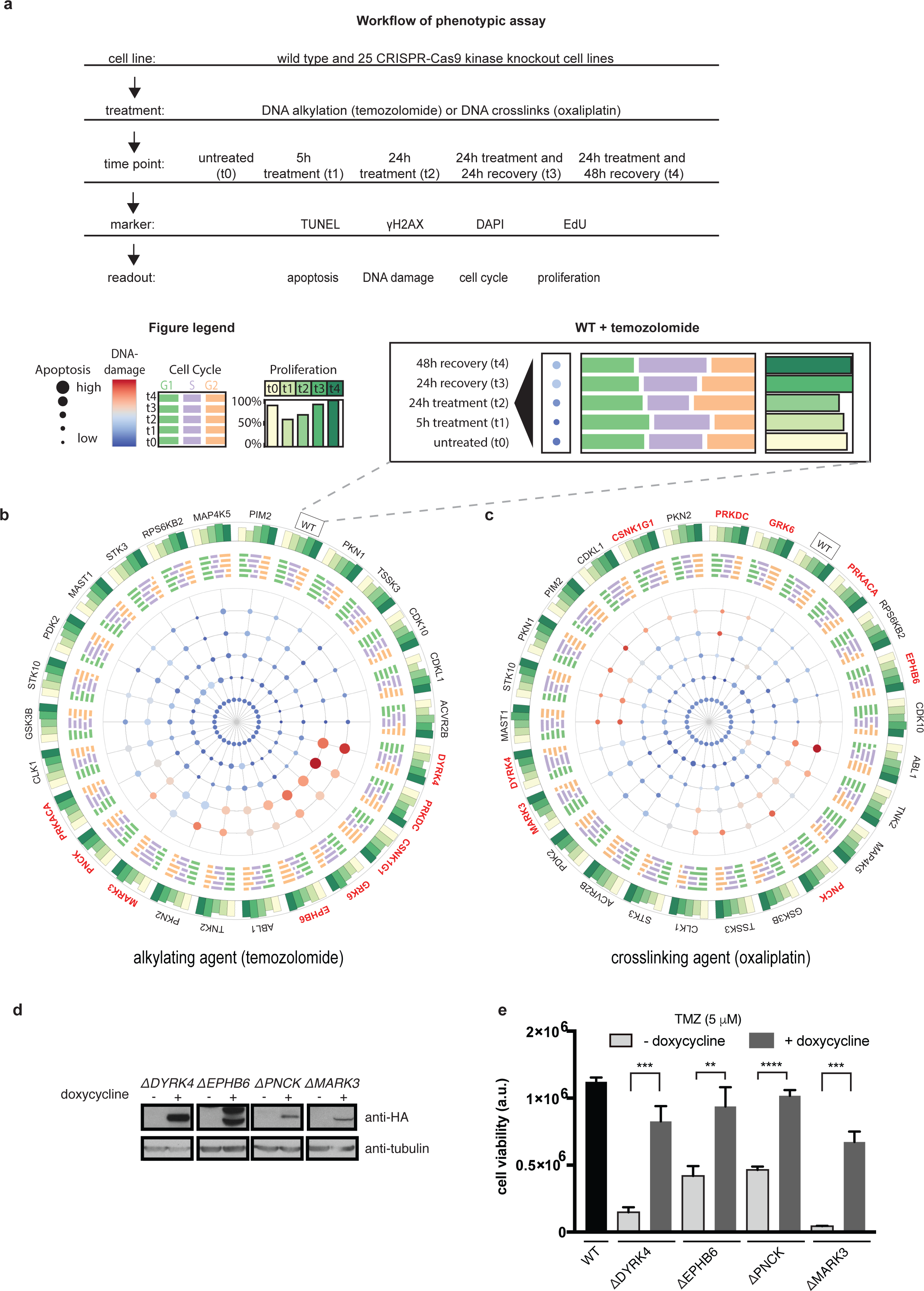
Phenotypic and genetic validation of sensitivity to alkylating agents. (a) Workflow of phenotypic screen. Wild-type and 25 CRISPR-Cas9 knockout cell lines were treated with alkylating or crosslinking agents (temozolomide or oxaliplatin, respectively) for 5 or 24 hours, after which drug medium was replaced with fresh medium to allow cells to recover from damage. EdU incorporation was performed for 40 minutes before harvest. Cells were fixed and co-stained using the following markers: TUNEL for apoptosis, anti-γH2AX for DNA damage, DAPI for cell cycle and EdU stain for proliferation. Cells were analysed by flow cytometry. For each stain, the six concentrations for each time point were summarized with an area under the curve (AUC) calculation. Bottom: Figure legend and phenotypic plot of wild-type HAP1 cells after temozolomide treatment (right). (b-c) Phenotypic plot for cell lines after alkylation-induced damage by temozolomide (b) or crosslinking induced damage by oxaliplatin (c). The inner most circle of the phenotypic plot shows DNA damage (color gradient of bubbles) and apoptosis (size of bubbles) for 4 different time points, t1 - t4, from center to periphery, and indicated cell lines. The next circle (middle) shows cell cycle distribution of cell lines in G1 (green), S (purple), and G2/M (orange) for time points t1 - t4, from center to periphery. The outermost circle shows proliferation changes of cell lines at time point t1 - t4. Cell lines are ordered hierarchically according to apoptosis and names are indicated outside of the phenotypic plot. Cells identified to be sensitive to carmustine (red names) show a high levels of apoptosis to temozolomide (7/10) but not to oxaliplatin (2/10), indicating that the survival response is primarily due to alkylating and not crosslinking lesions. Wild-type HAP1 cells (WT) are highlighted in a box. Figure legend as in (a). (d-e) Rescue of sensitivity to alkylating agents after reconstitution of knockout cell lines with HA-tagged, doxycycline inducible DYRK4, EPHB6, PNCK or MARK3 proteins. (d) Anti-HA immunoblot of the indicated cells lines reconstituted with the relevant cDNA after doxycycline induction. EPHB6, shows a characteristic smear of a fragmented membrane protein by immunoblotting. * indicates non- specific band. Tubulin was used as a loading control. (e) Upon temozolomide (TMZ) treatment, sensitivities of DYRK4, EPHB6, PNCK and MARK3 deficient cells are rescued after expression of exogenous proteins by doxycycline induction. Results are means of 3 replicates with standard deviations.

DNA damage as measured by γH2AX can either be a cause or consequence of apoptosis^37^. For a gene to be involved in the signaling or repair of DNA damage, we would expect to see higher levels of γH2AX preceding or coinciding with higher levels of apoptosis. For instance, DYRK4-deficient cells showed a peak of apoptosis 24 hours after treatment followed by a peak of γH2AX at 48 hours of treatment (Figure 2b). Hence, the γH2AX signal at the 48 hour time point may therefore be a consequence of apoptosis in DYRK4-deficient cells. This may also be the case for PNCK and PKN2 (Figure 2b). In contrast, PRKDC-deficient cells, which have a deficiency in DNA repair, showed the maximum levels of γH2AX and apoptosis early, at 24 hours after treatment, followed by a slow but coinciding recovery of γH2AX and apoptosis at 48 hours after treatment (Figure 2b). A similar, albeit weaker phenotype can be observed in cells deficient for CSNK1G1, EPHB6, MARK3, PRKACA (Figure 2b). Interestingly, we also observed a strong and persistent G2/M arrest in cells deficient for CSNK1G1 or EPHB6 (Figure 2b). Although the sensitivity of these kinase-deficient cells to alkylating agents revealed in our assay is as of yet unreported, it is in line with what is known about their function: CSNK1G1 has previously been reported to regulate the kinase CHK1, which is a cell cycle regulator following DNA damage^38^ whereas EPHB6 has been linked to the regulation of NPAT, a DNA damage signaling and cell cycle regulator^39^.

After confirming selected cellular survival phenotypes in our phenotypic screen, we next sought to validate gene-drug interactions in knockout cell lines by reconstitution of the wild-type genes. We selected the understudied kinases, DYRK4, EPHB6, PNCK and MARK3, as well as control kinases with resistance phenotypes, ABL1 and TSSK3 (Figure 1h,Supp. Figure 2a). After expression of HA-tagged inducible proteins (Figure 2d) in the deficient cell lines, we assessed whether cellular survival to DNA damage was reverted. Indeed, the sensitivity and resistance phenotypes could be corrected by recombinant expression of the relevant proteins hence establishing a coherent genotype-phenotype relationship (Figure 2e,Supp. Figure 2c-e). ABL1 and TSSK3 deficient cells, which showed resistance to doxorubicin or hydroxyurea in the survival screen, became significantly sensitive after reconstitution (Supp. Figure 2c-e), whereas DYRK4, EPHB6, PNCK and MARK3 which showed sensitivities in survival after alkylation-induced damage, became significantly more resistant after reconstitution of the respective wild-type genes (Figure 2e).

## Discussion

Unperturbed signaling of DNA damage is essential in guarding the genome against cancer^40^. At the same time targeting the DNA damage response has proven to be a successful strategy in cancer therapy. In this study, we have shown the response of 313 cell lines, lacking kinases involved in different cellular signaling pathways, against 10 diverse DNA damaging agents, including 7 commonly used chemotherapeutics. In doing so, we have identified unreported synthetic lethal and resistance gene-drug interactions. Moreover, we reveal that a sensitivity to carmustine may be important for a cluster of genes associated with chromatin accessibility. For selected cell lines, we further probe the synthetic lethality with carmustine by designing a phenotypic assay to investigate and validate gene-drug interactions in a broad manner. We show apoptosis, cell cycle, DNA damage and proliferation after alkylation or crosslink-induced damage for those cell lines. Moreover, we rescue the survival phenotype of DYRK4, EPHB6, MARK3, PNCK as a proof of principle for our study in reconstitution experiments. Our data suggests that some cancers with inactivated DYRK4, EPHB6, MARK3 or PNCK could be particularly vulnerable to alkylating agents. For example, EPHB6 is found to be downregulated in diverse metastatic cancers, including lung^41^, breast^42^ and brain^43^ cancers. Treatment of EPHB6 deficient cancers with the chemotherapeutic agents carmustine or TMZ may therefore represent a promising therapeutic strategy.

## Experimental Procedures

Experimental procedures can be found in the Supplemental Information.

## Author Contributions

M.O., P.B., MW and C.-H.L. carried out the experimental work. M.O., P.B., A.M., A.J., M.C. and J.F.daS. performed data analysis. Writing was by M.O. with J.I.L. The project was conceptualized by M.O. and J.I.L. Project administration was by S.K., J.M., F.C. and J.I.L. Funding acquisition was by S.K., F.C., J.M. and J.I.L. All authors reviewed and commented on the manuscript.

## Acknowledgments

Research in the Kubicek lab is supported by the Austrian Federal Ministry of Science, Research and Economy, the National Foundation for Research, Technology, and Development. M.O., P.B. and J.F.daS. were supported by FWF grant awarded to J.L. (29555 and 29763).

## Supplemental Experimental Procedures

**Supp. Figure 1.**
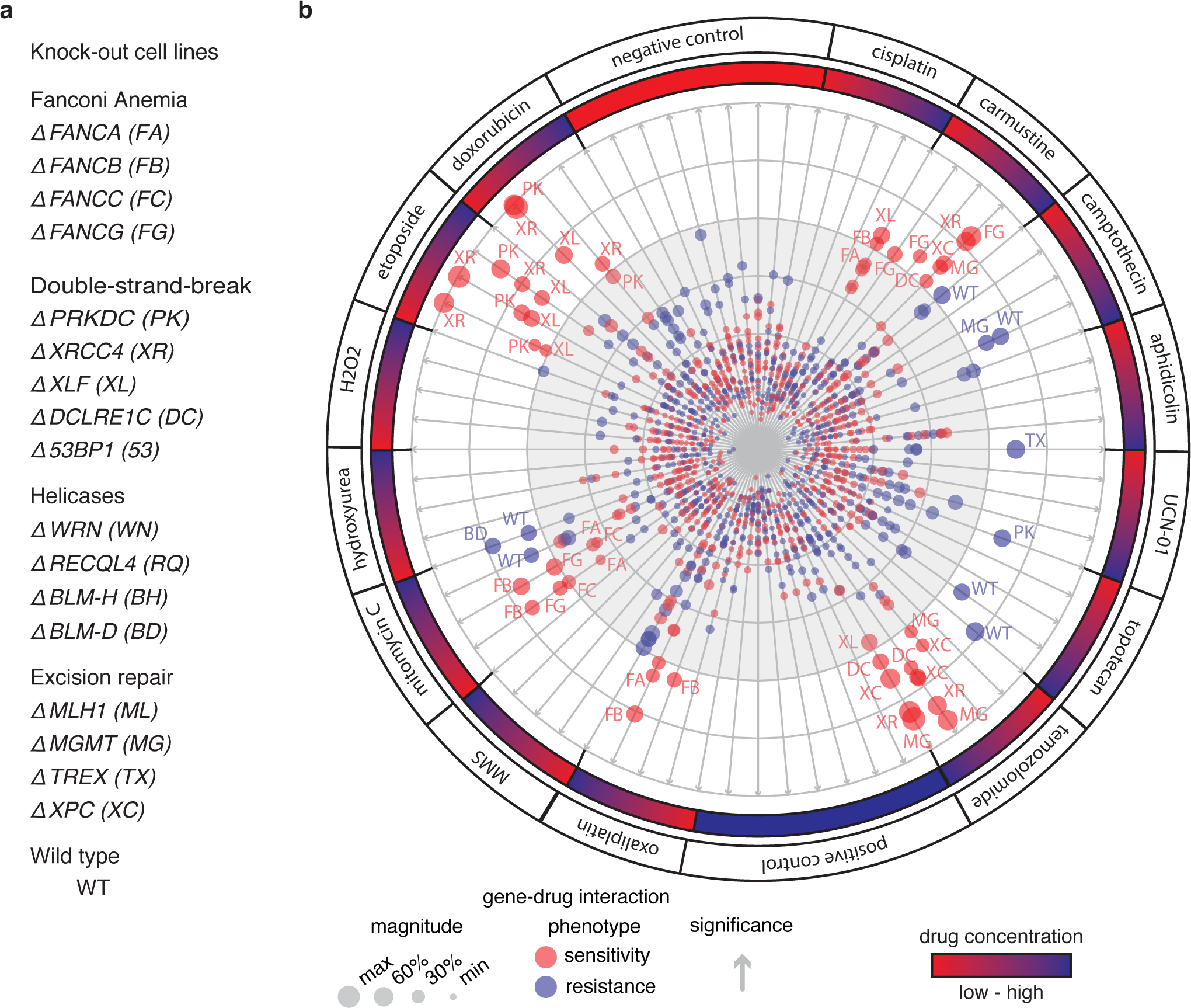
Survival of DNA repair deficient cell lines in response to DNA damage. (a) List of DNA repair deficient cell lines generated by CRISPR-Cas9 in human HAP1 cells (b) and their survival response to DNA damaging compounds are depicted in a circular bubble plot: DNA double-strand break repair deficient ΔPRKDC, ΔXRCC4, ΔXLF and ΔDCLRE1C are sensitive to DNA double-strand break inducing agents, doxorubicin and etoposide. Crosslinking repair deficient ΔFANCA, ΔFANCB, ΔFANCC and ΔFANCG are sensitive to crosslinking agents MMC, cisplatin and oxaliplatin. ΔMGMT and ΔXPC, deficient in the repair of alkylating adducts, are sensitive to the nonfunctional alkylating agent TMZ. Cell lines deficient in the repair of alkylation damage (ΔMGMT and ΔXPC) or crosslink induced lesions (ΔFANCG) are sensitive to the bifunctional alkylating agent carmustine. Each compound is represented by 4 dose points with a mean of 4 replicates per dose point. H_2_O_2_, Hydrogen peroxide; MMS, Methyl methanesulfonate; Negative control, DMSO; Positive control, 25X camptothecin (cytotoxic concentration). The color represents the drug-gene interaction, distance from the center indicates score/p-value and bubble size corresponds to the magnitude of the measured effect (over all other perturbations)^44^.

**Supp. Figure 2.**
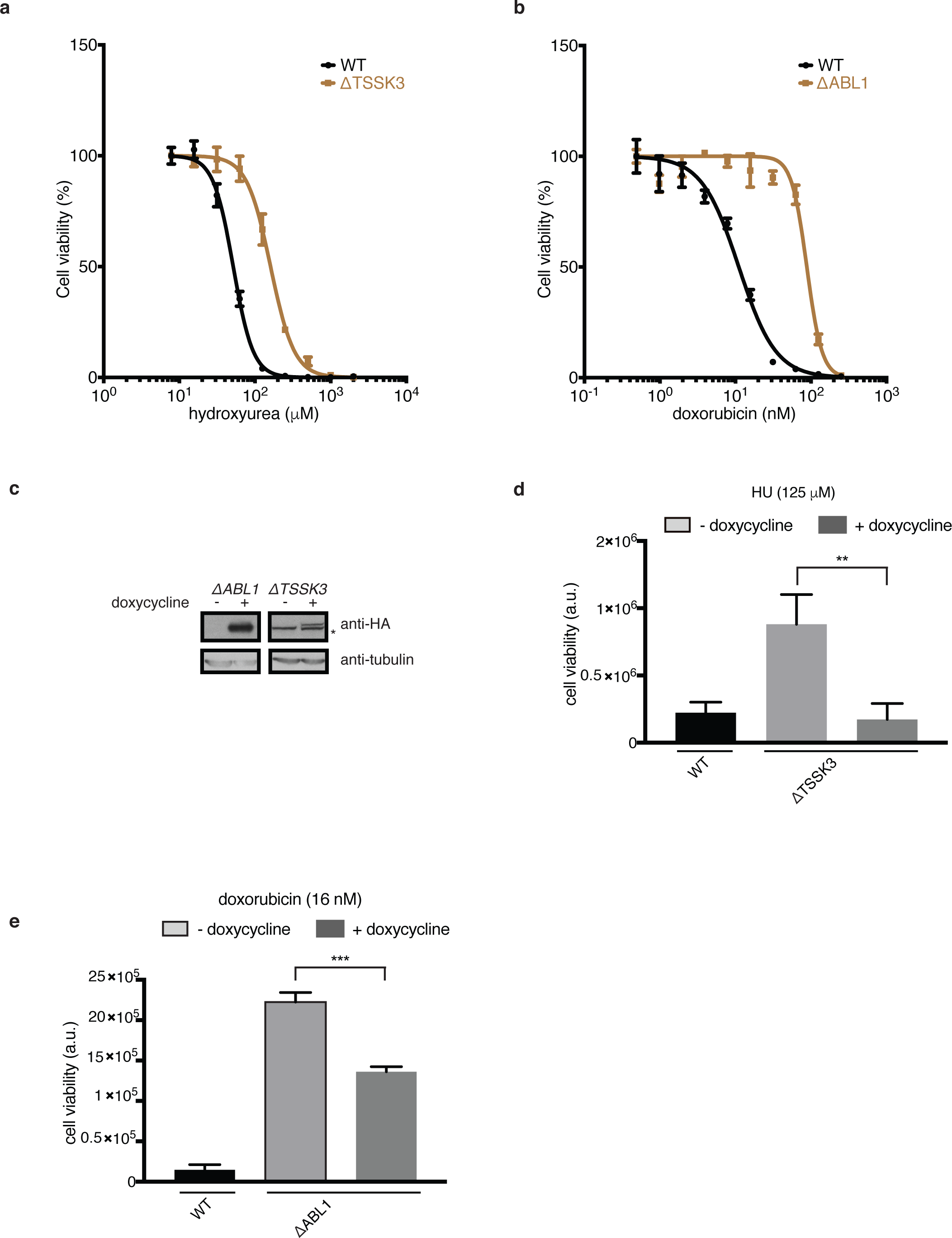
Rescue of resistance phenotypes after reconstitution of knockout cell lines with HA-tagged, doxycycline inducible proteins. (a-b) Validation of resistant survival responses of ΔTSSK3 to hydroxyurea and ΔBL1 to doxorubicin. (c) Anti-HA immunoblot of the indicated cell lines after doxycycline induction of the relevant cDNA. Expressed genes correspond to expected sizes. Wild-type control, WT, shows no affinity to anti-HA. * indicates non-specific band. Tubulin was used as a loading control. (d) Upon hydroxyurea (HU) treatment, resistance of TSSK3 deficient cells is rescued after expression of exogenous protein by doxycycline induction. Results are means of 3 replicates with standard deviations. (e) Setup as in (c) using resistant ALB1 deficient cells after doxorubicin treatment.

### Cell Culture

All HAP1 cell lines used in this work were generated using CRISPR-Cas9 gene editing technology in collaboration with Horizon Genomics (Vienna Austria) as single clones. They were grown in Iscove’s Modified Dulbecco’s Medium (IMDM) from GIBCO®, containing L-Glutamine and 25 mM HEPES and supplemented with 10% heat-inactivated fetal bovine serum (FBS) and 1% penicillin-streptomycin (P/S at 100µg/mL) and passaged according to standard protocol with trypsin and PBS. All cell lines were grown at 37°C in a 3% oxygen and 5% CO_2_-humidified incubator.

HEK293T cells used for virus production were expanded in Dulbecco’s Modified Eagle Medium (DMEM), supplemented with 10% FBS and 1% P/S. Cells were grown at 37°C in a 3% oxygen and 5% CO_2_ incubator.

### Knockout Confirmation by Sanger Sequencing

HAP1 kinase deficient knockout cell lines were validated for gene editing leading to a frame- shift mutation in the respective genes. We designed forward and reverse primers for each gene and purchased oligonucleotides from Sigma-Aldrich. For genomic DNA extraction, cells were treated with trypsin and washed twice with PBS, then resuspended in 100µL Direct PCR-Cell lysis solution with 2µL Proteinase K (20mg/mL). Wells were sealed and heated for 2.5 hours at 56°C, then 45 minutes at 80°C to inactivate Proteinase K, followed by PCR amplification. PCR amplification conditions: heat lid 110°C; 94°C 2 minutes; loop 35x (94°C 30 seconds; 337 55°C 30 seconds; 68°C 1 minute) 68°C 7 minutes. Then the PCR product was purified using Rapid PCR Cleanup Enzyme Set from BioLabs Inc., diluted 1:2 with double distilled water (ddH_2_O) and sequenced by Microsynth AG. Results were aligned to respective genes using Basic Local Alignment Search Tool, BLAST provided by NCBI.

### High-throughput Drug Screen

Indicated volumes and concentrations of compounds (**Supp. Table 3**) per well were transferred into 384-well plates (Corning 3712) from DMSO stock plates using acoustic transfer (Labcyte Echo 520). Wild-type or knockout HAP1 cells (at an amount of 1,000 cells) were seeded in 50 µl media into the compound-containing plates. Three days later cell viability was determined using Cell Titer-Glo (Promega). Compounds were used at 4 dose points with 4 replicates. For data analysis, the percentage of control was calculated and the signal of the DMSO treated sample was used to set values to 100% survival, while the 25X camptothecin (cytotoxic concentrations) signal was used to set the values to 0%. Survival circos plot with DNA repair deficient cells was created using TOPS^44^ for analysis and basic visualization. For kinome surival to DNA damage, areas under the curve (AUC) were calculated as a cumulative measure of compound potency by taking the sum of the mean of subsequent concentration points (as applied previously^45^).

### FACS Screen

Cell Culture: HAP1 cells were cultured as described above for 7 days until 80-90% confluence. Every HAP1 cell line was seeded into 4 Costar® 6-well cell culture plates, each plate at a density (cell number/well) according to the time-points of harvest: 160,000 for 5- hours treatment, 80,000 for 24 hours treatment, 20,000 for 24 hours post treatment and 10,000 for 48 hours post treatment.

Drug treatment: The day after seeding, cells were treated with 6 concentrations of the respective compound. The highest concentration- temozolomide, 250 µM; oxaliplatin, 780 nM- was chosen to moderately affect wild-type cells (10 - 30 % cell death). The compounds were serial diluted 1:2 from the highest to lowest dose. For the untreated control we used DMSO at a concentration corresponding to the lowest compound dilution. After 24 hours treatment, media from the remaining time points was aspirated and replaced with 2 mL of fresh (drug-free) IMDM medium. 40 minutes before each harvest 5-ethynyl-2′-deoxyuridine (EdU) at a concentration of 10 µM was added to each well.

Cell harvest: Cells were washed with 400µL phosphate buffer saline (PBS) and detached with 500µL trypsin, collected with 1 ml of medium, transferred into 96-deep-well (2ml) plates and centrifuged at 2,000 rpm for 6 min. The supernatant was carefully discarded, cell pellets were washed with PBS and re-suspended in 100µL fixing solution, containing 4% para-formaldehyde (PFA) and 0.1% Triton X, transferred into V-bottom shaped 96-well plates, incubated at 4°C and then stained.

FACS staining: 96-well plates, containing fixed cells, were centrifuged at 1,200 rpm for 6 min then washed with 50µL PBS. Pellets were re-suspended in TUNEL staining solution (*In Situ* Cell Death Detection Kit, TMR red, Sigma Aldrich) containing anti-phospho-H2A.X (Ser139) i.e. γH2AX, clone JBW301 (1:500 dilution, Sigma Aldrich) and incubated for one hour in the dark at 37°C. Then the pellets were washed three times with PBS (with centrifugations at 1,200 rpm for 6 min) and re-suspended with Click-iT EdU Alexa Fluor 488 Flow Cytometry Assay Kit staining solution (Thermo Fisher Scientific) containing secondary antibody (1:500 dilution, Alexa Fluor® 647 conjugate, goat anti-Mouse IgG (H+L), Thermo Fisher Scientific) for detection of γH2AX and incubated for one hour in the dark at room temperature (RT). Subsequently pellets were washed three times with PBS and re-suspended in DAPI (Sigma Aldrich) solution (1:1,000 dilution) and kept dark. Samples were measured using the BD LSRFortessa cell analyzer machine and data was analyzed using FlowJo v10.3.

### FACS Analysis

For analysis, dead cells were discarded using forward and side scatter and next single cell populations were gated using DAPI width and DAPI area following the Abcam PI staining protocol. Gates for γH2AX and TUNEL were set for all concentrations according to the untreated (DMSO) control. For all time points, to take all drug concentrations into account, an area-under-the-curve (AUC) of all six dose points was calculated and compared to untreated controls. This data is visualized in the phenotypic FACS plot. The cell cycle phases were determined by gating G1- and G2-phase as well as S-phase of diploid populations using a DAPI against EdU plot. Proliferating cells were determined by setting a threshold for cells with positive EdU incorporation and EdU positive signals were plotted against untreated control cells.

### Reconstitution of Knockout Cell Lines

For reconstitution of the respective wild-type genes in knockout cell lines, we used Gateway-cloning compatible vector backbones containing the gene of interest and a spectinomycin resistance cassette, from Addgene. These plasmids of the kind “pDONR223-XXX” were a gift from William Hahn & David Root and are published in Nature: Johannessen et al (2010 Nov 24.). Bacterial DH5α were grown on agar plates containing spectinomycin, from which a single clone was picked and cultured in LB-media containing spectinomycin overnight at 37°C. Plasmids were purified using Quiagen MidiPrep Kit and the LR-reaction was performed according to the Gateway Technology protocol provided by Invitrogen. We transferred the cDNA of the gene of interest into the doxycycline inducible pLIX_402 entry vector for mammalian expression and lentivirus production containing an Amp^R^ cassette. The entry clones were transformed into Mg^2+^/Ca^2+^ competent DH5α strains, amplified in an overnight culture in LB-media containing ampicillin and plasmids were extracted using Qiagen MidiPrep Kit. Lentivirus particles were produced using following plasmids: CMV-GFP, VSV-G and dR8.91. Virus was harvested for two days in the mornings and evenings. Knockout cells were infected with virus particles containing the respective gene, selected for 2 days with puromycin at 2 µg/mL and cells were propagated for another 2 days.

### Dose Responses

Dose response curves for temozolomide, oxaliplatin, hydroxyurea and doxorubicin were generated by seeding cells in 96-well plates (1,000 cells/well). The next day, compounds were added at the indicated concentrations. Cells harboring the reconstituted gene of interest were additionally treated with doxycycline at 1µg/mL every day. Three days after drug treatment, cell viability was measured using the CellTiter-Glo assay protocol (Promega).

### Immunoblotting and Antibodies

Cells were harvested and then lysed with RIPA lysis buffer (NEB) supplemented with protease and phosphatase inhibitors from Sigma. Western blots were performed according to standard protocols. Protein samples were separated using NuPAGE™ 4-12% gradient Bis-Tris Protein Gels from Invitrogen and MOPS running buffer at 120 V for 2 hours running time. The separated proteins were then transferred onto nitrocellulose membranes. To prevent unspecific protein binding, membranes were treated with blocking solution (5% milk in TBST) for 1 hour and primary antibodies were added at 1:1,000 to the blocking solution and incubated overnight at 4°C. The next day, membranes were washed 3x with TBST and incubated with secondary antibodies at 1:5,000 in 5% milk/TBST solution. Then membranes were treated with immunoblotting developer solution (GSE) for 1 minute and imaged in the dark. The following antibodies were used: primary; Rabbit Anti-HA tag antibody - ChIP Grade (ab9110, Abcam), secondary; Goat Anti-Rabbit IgG, HRP-linked Antibody (#7074, Cell Signaling).

### Clustering and HiC analysis

Cells were clustered with agglomerative clustering using the R package ConsensusClusterPlus. Cells and compounds were randomly sub-sampled for 10,000 times and clustered using complete linkage to increase the robustness of clustering. The distance was calculated using the Pearson correlation. Hi-C values for all genes in each cluster were retrieved from the covariate matrix of MutSigCV v1.2.01^46^ and the corresponding distributions were compared with Wilcoxon rank-sum test.

### Gene ontology term enrichment analysis

Gene ontology (GO) term enrichments were calculated using Enricher, a comprehensive tool for gene set enrichment analysis^47^. P-values were calculated using a Fisher's exact test and corrected for multiple hypotheses using a cut-off of p-value < 0.05. In order to filter for redundant and unspecific GO terms, we first removed all GO terms that are annotated to more than 70 genes and further summarized terms based on their Resnik semantic similarity^48^ using the tool ReviGO^49^.

